# Programming cell growth into different cluster shapes using diffusible signals

**DOI:** 10.1101/2020.08.29.273607

**Authors:** Yipei Guo, Mor Nitzan, Michael P. Brenner

## Abstract

Recent advances in genetic engineering technologies has made it possible to construct artificial genetic circuits and use them to control how cells respond to their surroundings. This has been used to generate spatial patterns of differential gene expression. In addition to the spatial arrangement of different cell types, another important aspect of spatial structure lies in the overall shape of the group of cells. However, the question of how cells can be programmed, and how complex the rules need to be, to achieve a desired tissue morphology has received less attention. In this paper, we attempt to address these questions by developing a mathematical model to study how cells can use diffusion-mediated local rules to grow into clusters with different shapes. Within our model, cells are allowed to secrete diffusible chemicals which can either directly regulate the growth rate of cells (‘growth regulator’), or indirectly affect growth by changing the secretion rate or the effect of other growth regulators. We find that (1) a single growth inhibitor can be used to grow a rod-like structure, (2) multiple growth regulators are required to grow multiple protrusions, and (3) the length and shape of each protrusion can be controlled using growth-threshold regulators. Based on these regulatory schemes, we also postulate how the range of achievable structures scales with the number of signals: (A) the maximum possible number of protrusions increases exponentially with the number of growth inhibitors involved, and (B) to control the growth of each set of protrusions, it is necessary to have an independent threshold regulator. Together, these experimentally-testable findings illustrate how our approach can be used to guide the design of regulatory circuits for achieving a desired target structure.

## I. INTRODUCTION

A fundamental goal in synthetic biology is to understand how cells can be programmed to generate a desired spatial configuration. In order to achieve such a collective goal, individual cells must make appropriate local decisions depending on where they are in the cluster and the current global state of the system. However, cells do not have direct access to these quantities and can only sense their immediate surroundings. Global information that needs to be accessible for cellular decision making must therefore be encoded in their local environment.

Recent advances in genetic engineering technologies have made it possible to encode desired sets of rules within the genetic programs of cells, and these have been used to create distinct spatial patterns [1–5]. For example, the use of a synthetic notch receptor system to encode changes in expression levels of cadherin molecules (in ‘receiver cells’ engineered with receptors that trigger downstream cellular responses when activated) upon contact with another cell type (‘sender cells’ engineered with ligands on cell surface) led to self-organization of clusters with distinct spatial arrangements of different cell types [3, 6]. In addition to juxtacrine signaling i.e. signaling through direct cell-cell contact (ligand/receptor systems), cells can also be engineered to communicate via diffusible signals [4–6]. In particular, a graded pattern of signaling activity was obtained by culturing engineered Hedgehog-responding cells next to engineered Hedgehog-secreting cells [4], and Turing-like patterns were generated by reconstituting an activator-inhibitor circuit of two diffusible molecules [5].

In addition to spatial patterns within a cell cluster (i.e. spatial arrangement of cells with different gene expression profiles), another important aspect of spatial structure lies in the form of the overall tissue shape. Even though this has received less attention in the field of synthetic biology, synthetic circuits can potentially be engineered to produce other forms of physiological outputs such as the regulation of growth/proliferation and death rates [1, 7], which would enable cells to grow into different cluster shapes. Nevertheless, before any experimental attempts, it is useful to first ask what are the rules that would enable cells to grow into a desired shape.

The question of how tissue morphology emerges has been widely studied in many biological systems, and there are many possible strategies cells can adopt [8, 9]. The ability to respond to local external chemical environments is particularly important in many developmental processes, where some cells can secrete morphogens which diffuse in extracellular space to induce concentration-dependent responses in other cells [10, 11]. In response to their local environment, cells can move [12], rearrange among themselves [13–16], produce different levels of actin/myosin [17, 18], undergo oriented cell division [19, 20], vary their growth rates [17, 18, 20, 21], amongst others. The precise regulation of any specific biological phenomenon is typically very complex, involving a combination of these mechanisms [17–20] or many types of signals or even multiple cell types, each following a different set of rules [14, 18]. However, when trying to grow a certain target structure in the lab, it is desirable to achieve a minimal working model, or to work with the minimal number of components. This requires an understanding of what elements are necessary to achieve a desired structural phenotype, and more generally, how does developmental complexity scale with complexity of these elements, or the cell-to-cell interaction rules [22].

In this paper, we attempt to address these questions by developing a framework for studying how different cluster shapes can emerge from cells regulating their growth rates based on their local chemical environment. Within our model, cells are allowed to secrete diffusible chemicals which can either directly regulate the growth rate of cells (‘growth regulator’), or indirectly affect growth by changing the secretion rate or the effect of other growth regulators. We find that (1) a single growth inhibitor can be used to grow a rod-like structure, (2) multiple growth regulators are required to grow multiple protrusions, and (3) the length and shape of each protrusion can be controlled using growth-threshold regulators. With these regulatory schemes, we also illustrate how our approach can be used to infer how developmental complexity (i.e. the range of achievable structures) scales with model complexity (i.e. the number of signals) for any given initial cluster. In particular, we find that the maximum possible number of protrusions increases exponentially with the number of growth inhibitors involved, and to control the growth of each set of protrusions, it is necessary to have an independent threshold regulator.

## II. RESULTS

### A. Modeling diffusion-based morphogenesis

Suppose every cell has the potential to secrete *q* different chemicals, with the secretion rate 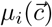 of chemical *i* potentially regulated by the local chemical environment 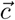 of the cell (Fig. 1):

**FIG. 1.**
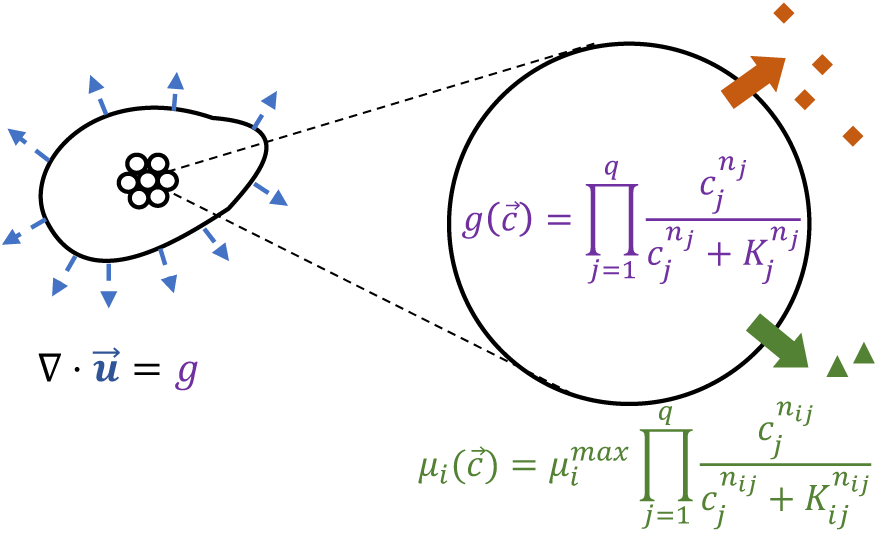
Schematic of the model. We consider a cluster of cells, each having the potential to secrete *q* different diffusible chemicals. The secretion rate 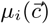 of each of these chemicals *i* = 1, …, *q*, and the growth rate 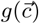 of each cell depend on the local external chemical environment 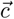 of the cell. Given the spatial profile of growth rate inside the cluster, we can then solve for the velocity of cell flow 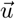, which specifies how the shape of the tissue changes over time (blue arrows).

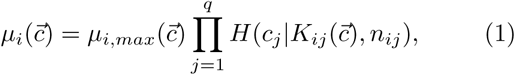

where *µ*_*i,max*_ is the maximum secretion rate of *i*, and *H*(*c*|*K, n*) = *c*^*n*^*/*(*K*^*n*^ + *c*^*n*^) is the Hill function.

We consider a 2D cluster of cells in a liquid medium and assume that the growth rate is much slower than the secretion, degradation and diffusion rates, such that the spatial profile of chemical concentrations satisfy the set of steady-state reaction-diffusion equations:

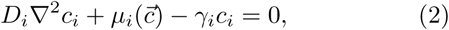

for *i* = 1, …, *q*, where *D*_*i*_ is the diffusion coefficient, *γ*_*i*_ is the degradation rate of *i*, and we assume the concentrations vanish far from the cell cluster.

We also allow the chemicals to be *growth regulators* that regulate the growth and division rate *g* of each cell (Fig. 1):

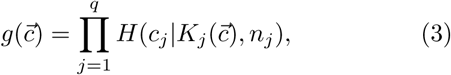

where we have taken the maximum growth rate to be 1. In both the secretion and growth regulatory functions (Eqns. 1, 3), we take the Hill coefficients *n*_*ij*_ and *n*_*j*_ to be either 0 (when *j* does not participate in the regulation), or very large in magnitude (negative when *j* is an inhibitor, positive when *j* is an activator) such that the Hill functions can be thought of as threshold functions.

Within a tissue, or a cellular cluster, the growth zone, which marks the region where cells can grow and divide, is determined by Eqns.1-3. To simulate the dynamics given the growth zone, we model the cell cluster as an incompressible cellular ‘fluid’ of constant density, such that the velocity **u** of cell flow is given by (Fig. 1):

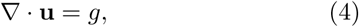

with **u** = −∇*P* and we impose the condition that the pressure is *P* = 0 at the cluster boundary. For any *g*(**x**), the velocity of the tissue along its boundary can then be obtained using boundary integral methods (Appendix A).

The equations above specify the dynamics for any regulatory mechanism, including the number of chemicals involved and the role of each of these chemicals i.e. how they affect growth and secretion of other chemicals.

Given a fixed set of growth rules, the resulting cell cluster will also depend on the initial configuration. In particular, with an initial circular cell cluster, if there are no spontaneous instabilities, concentration profiles and hence the cluster will remain circularly symmetric as it grows. For more complex shapes to emerge, there must be some symmetry breaking mechanism. In biological systems, the initial cluster shape is often determined by external conditions such as an external concentration gradient. Furthermore, during the early stages of development, the timing, order and plane of cell divisions are typically highly regulated and possibly governed by a separate set of rules encoded within the cell. When trying to grow these tissues in the lab, one can imagine creating molds or patterned environments for initializing the arrangement of cells. However, our goal here is not to explore what can be achieved throughout the whole space of initial conditions. Instead, we will illustrate how our framework can be used to inform how regulatory circuits should be designed in order to grow desired structural features from a given initial cluster shape.

Inspired by the example of developing bird beaks, which has been shown to be conic sections with a localized growth zone near the tip [23], we choose to initialize the system with a 2-dimensional parabolic cell cluster, and assume that because of an initial symmetry breaking event, only cells in the front half of the initial cluster (i.e. cells with *x*−coordinate greater than *L*_0_*/*2, where *x* = 0 corresponds to the vertical edge of the cluster and *x* = *L*_0_ is the location of the initial parabolic tip) can divide. Nevertheless, we find that an initial elliptic cell cluster gives rise to qualitatively similar structural features (Fig. S1).

In the rest of the paper, we will make use of concrete examples of this model to explore how the range of achievable structures varies with the type of regulatory mechanism and the number of signals involved in the growth regulation. The regulatory schemes we explore in this paper do not exhibit spontaneous instability, and we return to this point in the discussion section (Section III).

### B. Single growth inhibitor can give rise to an elongating rod-like structure

If all cells within the tissue were to grow uniformly at the same rate, the cluster will expand in all directions, maintaining its shape while it grows. Within our model, in order for the tissue to elongate preferentially in one direction, there must therefore be growth heterogeneity across the cluster, something that can only be achieved with a growth regulator.

We first consider the simplest scenario where all cells are secreting only a single growth inhibitor *X* at a constant rate *µ*_*X*_. The inhibitor concentration *c*_*X*_ satisfies the following non-dimensional equation:

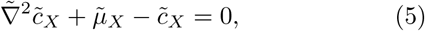

where the rescaled length scale 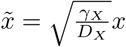, the rescaled inhibitor concentration 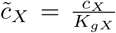 with *K*_*gX*_ being the threshold concentration above which cells stop growing (i.e. 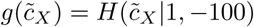 in Eqn. 3), and the effective secretion rate 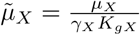.

Given any initial cluster size, the subsequent dynamics are therefore determined only by a single parameter 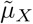, with the growth zone being the region where 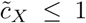 (Fig.2a). If 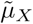 is too low, no cells will be inhibited; if 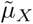 is too high, no cells can grow. For intermediate values of 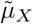, we find that the growth zone is localized near the tip of the parabolic cluster, and the size of the growth zone decreases with increasing 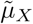 (Fig.2b). With the growth zone at the tip, the cluster grows a rod-like structure regardless of the size of the growth zone (Fig.2b). In the long time limit, the chemical environment at the tip and hence the size of the growth zone stays approximately constant during growth (Fig.2c,d), with this steady-state size also decreasing with 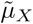 (Fig.2d).

**FIG. 2.**
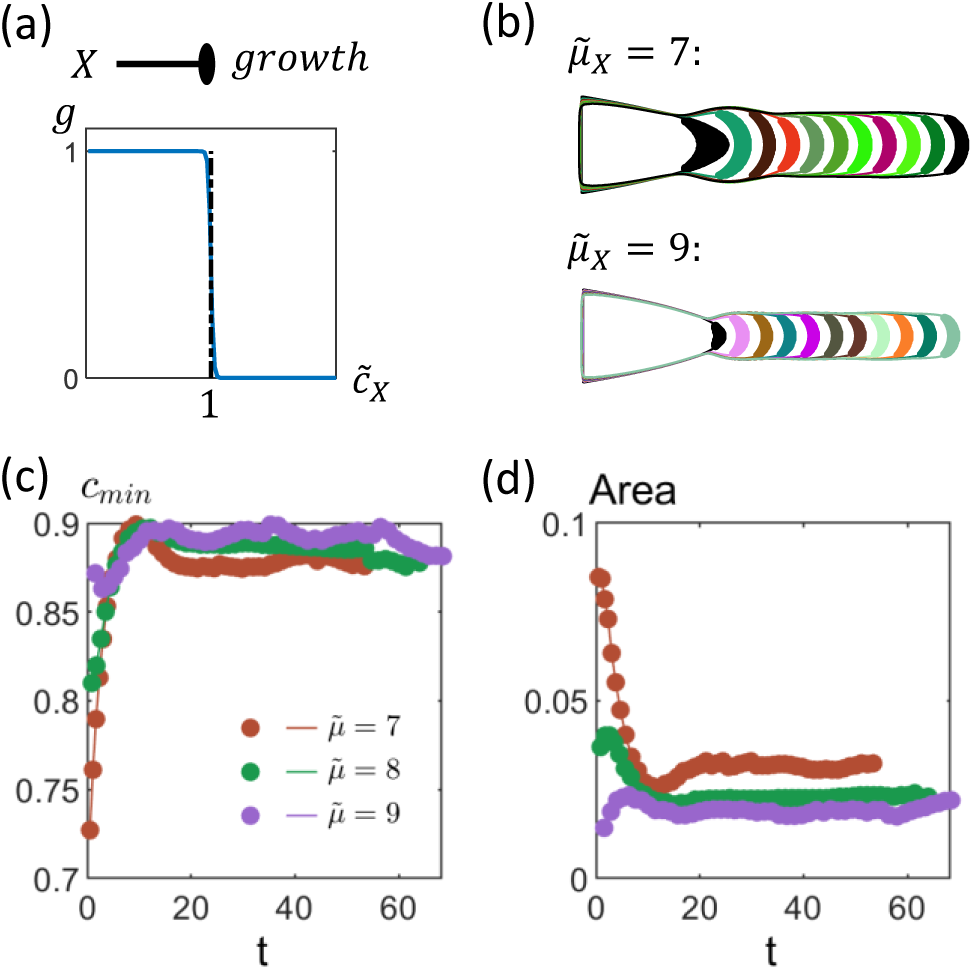
Dynamics of a parabolic 2-D cluster with cells secreting a single growth inhibitor. (a) We treat *X* is a growth inhibitor, such that cells can only grow if the concentration of *X* is below a threshold i.e. 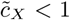. (b) Only cells at the tip of the tissue can grow, with the initial size of the growth zone (black region) decreasing with increasing effective secretion rate of inhibitor 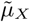. As cluster grows, a rod-like extension emerges. The different colored regions represent the growth zones at different times. (c) Rescaled inhibitor concentration 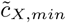 at the tip initially increases but reaches a steady-state level where it stays approximately constant. (d) Area of growth zone eventually stays approximately constant as the rod-like extension grows. The steady-state area decreases with 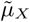. [Other parameters: initial tissue length 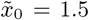, initial tissue width 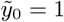]

Such a mechanism can therefore potentially be used for growing rod-like extensions, with the cluster stop growing when cells encounter depletion in nutrients, oxygen or other signals necessary for growth.

This result also implies that within our model and for an initial parabolic cell cluster, a single growth inhibitor alone cannot give rise to anything other than a single protrusion with approximately constant width. We therefore ask how more complex shapes can develop with the use of more chemical signals. In the rest of the paper, we will focus on two specific features: (a) the growth of multiple protrusions and (b) the control over how the width of a protrusion varies as it grows.

### C. Growing multiple protrusions with multiple growth inhibitors

In order for multiple protrusions to develop, there has to be multiple peaks in the normal velocities along the cluster boundary. One way for this to occur is for the tissue to have multiple growth zones - these do not necessarily need to be present at the initial state as long as there is potential for these distinct growth zones to emerge during growth.

Within the framework of our model, a chemical can either influence the secretion rate of other chemicals (Eqn. 1), affect growth rate regulation by other chemicals through the growth threshold or is itself a growth regulator (Eqn. 3). We find that since the concentration of growth regulators most directly determines whether a cell can grow, including additional growth regulators can change growth zone shapes most drastically. Nevertheless, for each additional growth regulator to provide spatial information that the other growth regulators could not have provided, the secretion of these additional growth regulators must be regulated such that not all cells are producing the same set of growth regulators.

A simple mechanism for increasing the number of possible protrusions (through introducing the potential for growth zones to split) is to allow cells within a growth zone to produce an additional growth inhibitor. We will first illustrate how this works with two growth inhibitors, before generalizing to the scenario of having multiple growth inhibitors.

#### It is possible to grow up to three protrusions with just two growth inhibitors

We consider here the case where *X*, in addition to being a growth inhibitor, also inhibits the secretion of a second growth inhibitor *Y* (Fig. 3a). This implies that *Y* is only produced when *X* is below some threshold level *c*_*X*_ *< K*_*s*_ (Fig. 3a), i.e.

**FIG. 3.**
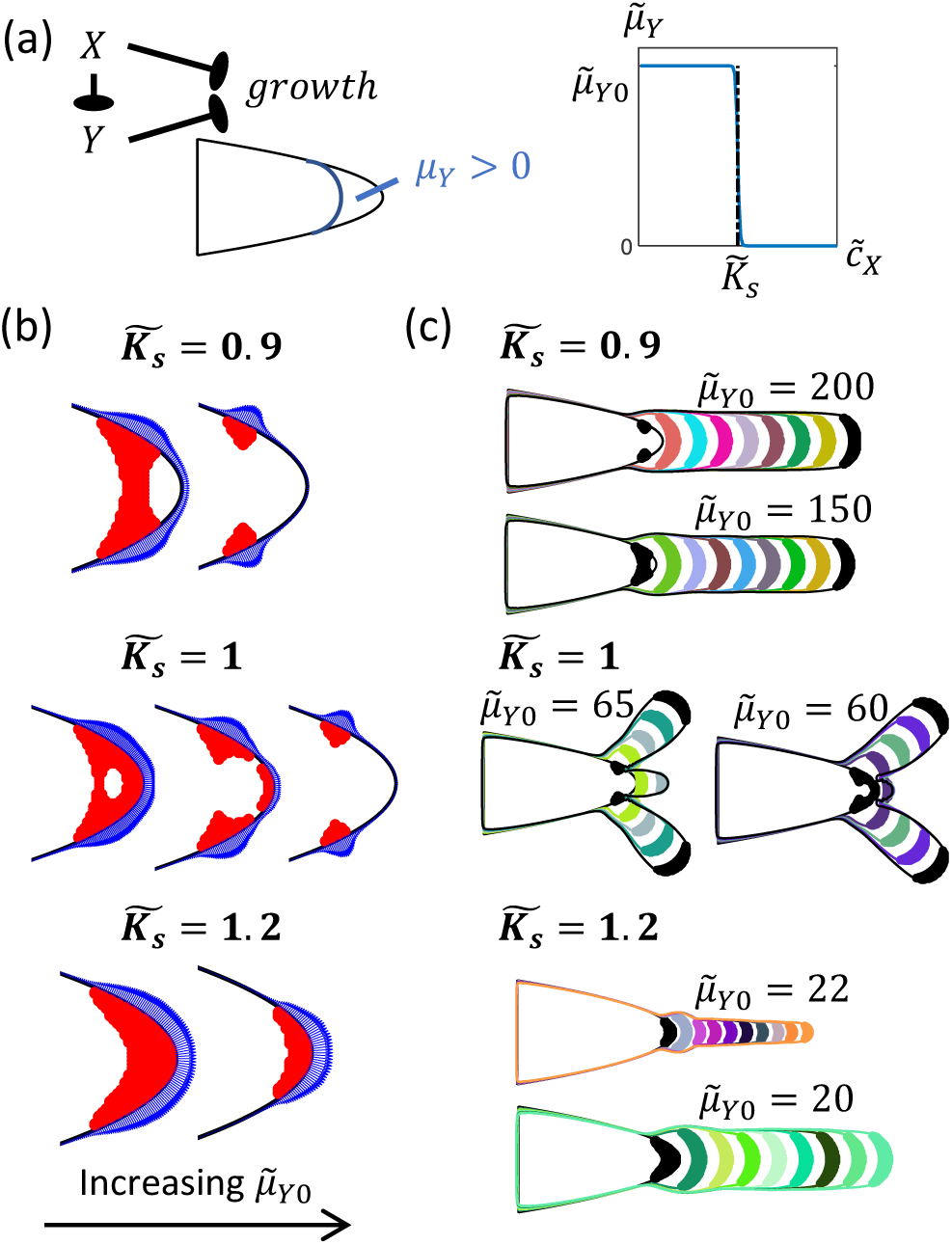
Dynamics of cluster when cells secrete two growth inhibitors. (a) We consider here the case where both *X* and *Y* inhibits growth. In addition, *X* also inhibits the production of *Y*, such that its secretion rate 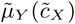 is a threshold function with *Y* only produced near the tip of the cluster where 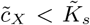. (b) The growth zone (in red) takes on different shapes depending on the values of 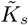 and 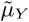. When 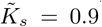, the region of the growth zone closer to the tissue tip stops growing first as 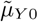 is increased (top row, left: 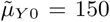, right: 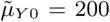). When 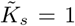, growth zone depletes from its center as 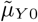 increases (middle row, left: 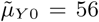, middle: 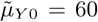, right: 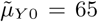). When the ratio of secretion threshold to growth threshold 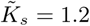, the growth zone shrinks from the left boundary as 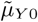 increases (bottom row, left: 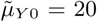, right: 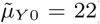). The growth zone therefore remains attached to the tip of the tissue. The blue regions consist of blue arrows perpendicular to the tissue boundary, with the length of the arrows proportional to the boundary velocity at that point. (c) Cluster dynamics for different secretion thresholds 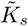, with the different colored regions representing the growth zones at different time points. Cluster grows multiple protrusions when 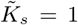. There is only a single growth zone and hence a single protrusion when 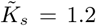. When 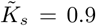, the cluster grows a single rod-like protrusion even though there are initially multiple growth regions. [Other dimensionless parameters: 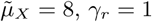.]

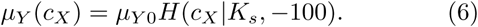

The steady-state concentrations therefore satisfy the following set of non-dimensional equations:

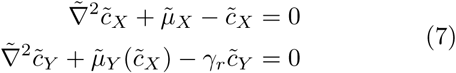

where the rescaled length scale 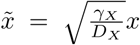, rescaled concentrations 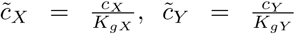 and 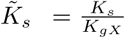, effective secretion rates 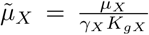 and 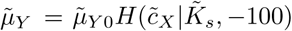 with 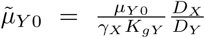, and the rescaled *Y* degradation rate 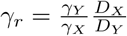.

The corresponding growth condition is given by:

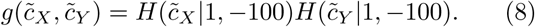

Fixing the secretion rate of *X* to be 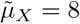, we find the inclusion of *Y* allows the growth zone to take on different shapes depending on the values of 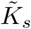 and 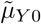 (Fig.3b). In the low 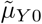 limit, the growth zone is determined solely by *c*_*X*_. As 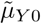 increases, some of the cells that were not inhibited by *X* may now be inhibited by *Y*. This reduces the size of the growth zone. However, the way in which the shape of the growth zone changes with increasing 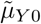 depends on the secretion threshold 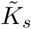 (Fig.3b). This is because the growth zone starts being depleted at the point where *c*_*Y*_ is the highest,

In particular, if cells secrete *Y* only when their growth is inhibited by 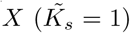, the growth zone is depleted from the center of the original growth zone (Fig.3b). The presence of this ‘hole’ changes the velocity profiles along the boundary, leading to multiple peaks in velocity along the boundary. This in turn gives rise to a pair of protrusions extending from opposite sides of the tissue, in addition to the middle tip protrusion. The tip protrusion eventually stops growing as *c*_*X*_ (which includes contributions from cells in both side protrusions) increases. Depending on the value of 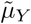, which determines the initial fraction of growing cells and hence how fast *c*_*X*_ increases, the tip protrusion can reach different lengths before it stops growing (Fig. 3c). However, the side protrusions continue to grow because the increase in number of cells (and hence total production of *X*) is offset by these protrusions getting further away from the main bulk of the cluster.

This mechanism shows that having two growth inhibitors can give rise to a maximum of 3 protrusions. This is nevertheless only an upper bound. If 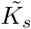 is sufficiently large such that the growth zone depletes from the boundary closer to the bulk of the tissue (Fig.3b), there will be just a single growth zone and hence a single protrusion (Fig. 3c). Furthermore, even with multiple distinct growth zones, it is possible for the cluster to grow a single rod-like structure similar to when cells only produce *X* (Fig.3c).

#### The potential number of protrusions scales exponentially with the number of growth regulators

This same strategy can be repeated with more than 2 growth inhibitors. If for example a third growth inhibitor *Z* is produced within the existing growth zone (i.e. when both *X* and *Y* levels are low), this can again split the existing growth zones, doubling the number of potential protrusions. This argument therefore implies that with *q* ≥ 2 growth regulators, it is possible to get a maximum of *z*_*max*_ = 3 × 2^*q*−2^ protrusions. Equivalently, *q*_*min*_ = log_2_(*z/*3) + 2 is the minimum number of growth regulators we would need to get *z* protrusions. Together, the potential number of protrusions scales exponentially with the number of growth regulators.

### D. Regulating the growth of individual protrusions using threshold-regulators

In addition to growing multiple protrusions, one may also wish to regulate the growth of each of them i.e. control how the width changes over time and for the cluster to stop growing by itself. We first ask how this can be achieved for a single protrusion, before discussing the general case of controlling multiple protrusions.

#### A growth threshold regulator, together with a growth inhibitor, can give rise to a cone-like protrusion

We saw previously that with a single growth inhibitor, the protrusion will grow with approximately constant width. This implies that for its width to change over time, additional chemicals are required. A mechanism for the protrusion to grow a sharp tip is for the growth zone to shrink and the center of the growth zone to shift closer to the tip as the protrusion grows. This is in fact what happens in the development of bird beaks [23].

Inspired by this, we ask here how such a phenomenon can arise. One way this could happen is if the growth threshold *K*_*gX*_ of the growth inhibitor *X* decreases over time. However, since any growth rule must be local, the time dependence of *K*_*gX*_ must occur through the dependence on the chemical environment i.e. 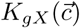. We therefore considered the possibility of having a second chemical *Y* that reduces *K*_*gX*_. For simplicity, we chose a linear function for this threshold regulation (Fig.4a):

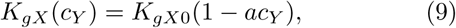

where *K*_*gX*0_ is the baseline growth threshold when *Y* is absent, and *a* controls how strongly *Y* regulates *K*_*gX*_.

Nevertheless, we find that the secretion of this threshold regulator is not a sufficient condition for the growth zone to decrease in size − it is necessary for the secretion of *Y* to be regulated such that only certain regions of the tissue are secreting *Y*. This is because if *Y* is secreted at a constant rate by all cells, *c*_*Y*_ at the tip will not increase as the tissue grows (just like for the growth inhibitor *X*). In this case, cells at the growing tip does not have any information of how far they are from the bulk of the tissue, and the cluster again grows a rod-like structure (Fig. 4b).

**FIG. 4.**
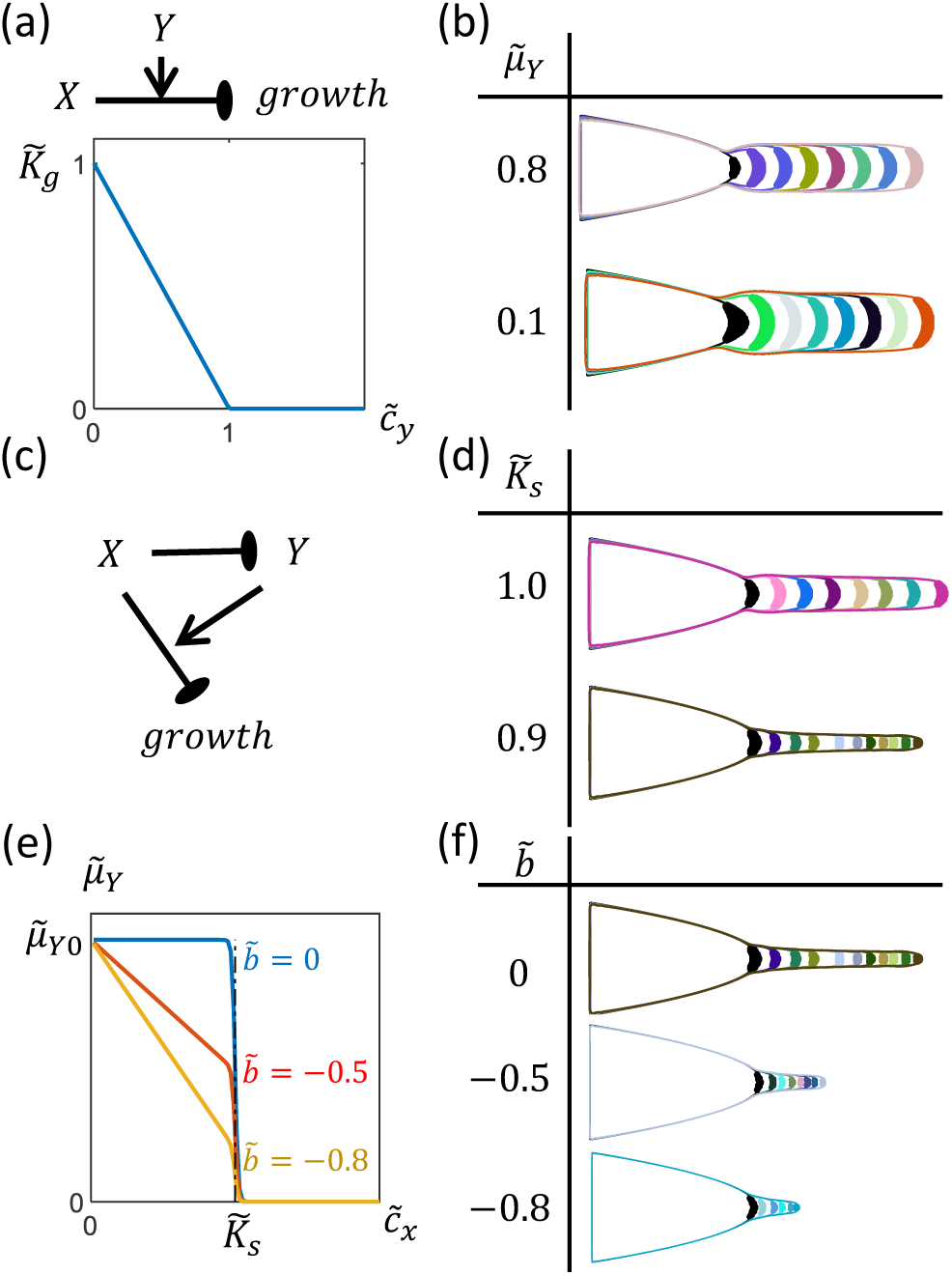
Dynamics of cluster when cells secrete a growth threshold regulator in addition to a growth inhibitor. (a) *Y* reduces the growth threshold of *X*. Here, we have chosen the rescaled growth threshold 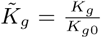 to decrease linearly with the rescaled concentration 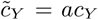. (Eqn.9) (b) The cluster grows a rod-like structure when all cells secrete *Y* at a constant effective rate 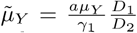. (c) We consider the scenario where *X* inhibits the production of *Y* such that only a region at the tip of the tissue can secrete *Y*. (d) For the scenario described in (c), it is possible for the protrusion to become narrower over time if the secretion threshold is less than the growth threshold 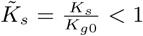 (top: 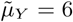, bottom: 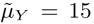). (e) We also allow the maximum secretion rate of *Y* to decrease linearly with *c*_*X*_. (Eqn.10). (f) When 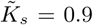, a stronger regulation of *µ*_*Y,max*_ (more negative 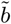) produces sharper cone-like structures (top: 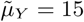, middle: 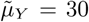, bottom: 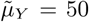).[Other dimensionless parameters: 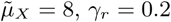

In order for only certain regions of tissue to secrete *Y*, we allow the secretion rate of *Y* to depend on the concentration of *X* as in the case of 2 growth inhibitors (Eqn. 6). The steady-state diffusion-reaction equations for this context are the same as if there were 2 growth inhibitors (Eqn.7), except that the rescaled concentrations are now 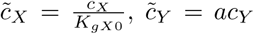 and 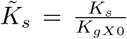, the effective secretion rates 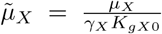 and 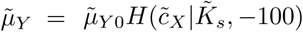 with 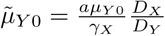.

For the growth zone to decrease in size, *c*_*Y*_ at the tip needs to increase as the tissue grows. We find that an effective way for this to occur is for cells to secrete *Y* only when 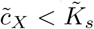 (Fig. 4c), and for 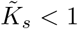 such that only a subset of growing cells (close to the tip of the tissue) are producing *Y* (Fig. 4d). As the tissue grows, the secretion region of *Y* increases and hence the maximum value of *c*_*Y*_ increases. This increase in *c*_*Y*_ reduces the size of the growth zone over time, giving rise to a narrowing of the protrusion (Fig.4d). The protrusion eventually stops growing when the the area of the growth zone vanishes.

The length of the cone-like protrusion can be controlled by regulating the rate at which the size of the growth zone decreases. We find that this can be achieved through regulating the maximum secretion rate *µ*_*Y,max*_ of *Y* in the regime where *Y* is being secreted (i.e. 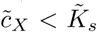). More specifically, we allow *µ*_*Y,max*_ to increase with decreasing *c*_*X*_ (Fig. 4e):

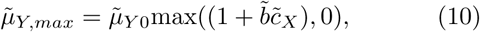

where as before the overhead ∼ indicates the corresponding rescaled variables, and 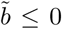 is a dimensionless parameter that controls how strongly *X* regulates *µ*_*Y,max*_. We find that a stronger regulation of *µ*_*Y,max*_ can give sharper cones (Fig. 4f). This is because as the protrusion grows, *c*_*X*_ decreases. A more negative *b* therefore results in a greater increase in the production of *Y*, and hence a faster shrinkage of the growth zone.

#### An independent threshold-regulator should be used to control the growth of each set of protrusions

If there are multiple protrusions such as in the presence of multiple growth inhibitors, we would expect that having a single growth threshold regulator would affect all protrusions in a similar or correlated way. Therefore, to control the lengths of these protrusions independently, we should have a different growth-threshold regulator for each set of protrusions. This set of growth-threshold regulators can regulate the same growth inhibitor, but their secretion rates (including the condition under which they are secreted) and their effect on the growth threshold may vary. Therefore, each regulator may be active only in the chemical environment of the protrusion it controls. Note however that due to the parabolic symmetry in our problem, the side protrusions can only be controlled in pairs.

## III. DISCUSSION

There are many possible ways by which tissues can change their shape during the developmental process. In the context of tissue elongation [24], this can occur through localized proliferation such as in bird beaks [23], oriented cell divisions such as in the *Drosophila* wing disc epithelium [20], cell intercalation [15, 16, 25], and elongation of individual cells [24]. All of these mechanisms fundamentally require cells to sense their local environment and respond by varying gene expression levels, which in turn regulate the growth rate of cells, the production rate of various intracellular proteins, the secretion rate of diffusible signals, the polarization of cells, amongst others. How these responses can be programmed, and how complicated the regulatory mechanisms need to be, to achieve a desired tissue shape and structure is a fundamental question in biology.

Here we explore changes in shape that arise solely from differential growth rates across the tissue, and investigate this in the context of growth rate regulation via diffusible morphogens. By using a bottom-up approach, we find with an initial parabolic cell cluster that it is possible to grow a rod-like extension with just a single growth inhibitor, and that having multiple growth inhibitors allow for multiple protrusions while growth threshold regulators can be used to regulate the shape and length of each protrusion, allowing protrusions to be cone-shaped rather than rod-like. In addition to what is achievable with chemicals with different functions, the limits of each regulatory circuit can also be inferred from these results given a fixed initial condition. In our example, a single growth inhibitor alone (with or without growth threshold regulators) cannot give rise to multiple protrusions. More generally, with the regulatory schemes studied here, the maximum possible number of protrusions grows exponentially with the number of growth inhibitors involved, and there should be an additional growth-threshold regulator for each set of protrusions one wishes to control independently. These results provide a lower bound for the number of signals cells need to achieve a certain goal. Presumably similar analyses and general arguments can be made for other structural features and the corresponding regulatory mechanisms. Furthermore, there may be multiple sets of rules that could give rise to qualitatively similar structural features. Such an analysis can therefore also be useful for comparing different regulatory schemes in terms of their capacity to generate complex structures.

Turing-like instabilities have been used to explain pattern formation in many biological systems and can give rise to digit patterning [26]. Even though Turing patterns may be a convenient way of getting a large number of protrusions with very few chemicals, they typically operate over narrow parameter ranges, and the nature of the Turing mechanism suggests that the type and features of the patterns may change drastically as the tissue grows and changes its shape. Coupling such a mechanism to growth regulation therefore entails another degree of complexity that would probably require fine-tuning of the parameters. Here, we take a different approach and our results show that it is possible to obtain multiple protrusions with other regulatory schemes that do not involve such spontaneous instabilities.

Identifying possible regulatory mechanisms and the minimum number of signals needed for achieving a certain structural feature may provide insight into natural developmental systems, and is especially useful for engineering these clusters synthetically. Given the advancement in genetic engineering techniques, the concrete predictions we have could potentially be tested in the lab. More importantly, our framework can be used to guide the design of regulatory circuits for achieving a desired target structure. In particular, this model could be used to test if a proposed/hypothesized regulatory scheme can achieve a particular structure and if so, what are the relevant parameter regimes. Furthermore, even though we have assumed that all cells in the cluster follow the same set of rules, our framework may potentially be extended to include multiple cell types, with each cell type following a different growth rule, or other processes such as cell differentiation (where cells change from one type to another based on local rules) and cell reorganization driven by differential adhesion.

## MATERIALS AND METHODS

All simulations are carried out using custom code written in MATLAB (R2019a). These codes can be found in the GitHub repository: https://github.com/yipeiguo/ProgrammingGrowth

## ACKNOWLEDGMENTS

This research was funded by the National Science Foundation through DMS-1715477, MRSEC DMR1420570, and ONR N00014-17-1-3029. MPB is an investigator of the Simons Foundation. M.N. acknowledges support from James S. McDonnell Foundation, Schmidt Futures, Israel Council for Higher Education, and the John Harvard Distinguished Science Fellows Program within the FAS Division of Science of Harvard University.

The authors declare no competing interests.

## Appendix A: Solving for velocities at tissue boundary

The velocity of the tissue boundary given any spatial growth rate profile 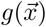 is found from solving the equation:

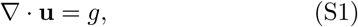

with **u** = −∇*P* and we impose the condition that the pressure is *P* = 0 at the cluster boundary.

To derive the boundary integral equation used for finding the boundary velocities, we follow closely the approach used in Ref. [27].

To deal with the boundary condition *P* = 0, we decompose the potential *P* into 2 components:

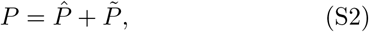

Where

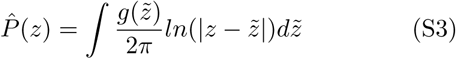

is the contribution from the growth rates of all cells within the tissue, while 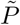 is the contribution due to the presence of the tissue boundary and satisfies:

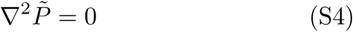

inside the tissue, with 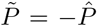 on the tissue boundary *∂*Ω.

The total cell velocity can therefore also be written as the sum of the two contributions: **u** = **û** + **ũ**, with 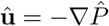 and 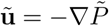. Since *P* is constant on *∂*Ω, the tangential component of the velocity *u*_*τ*_ = 0, and to solve for the evolution of the tissue shape, we are interested in finding the normal velocity component *u*_*n*_ = *ũ*_*n*_ + *û*_*n*_.

To do so, it is convenient to consider *∂*Ω as a contour of total length *L* in complex plane, and parameterise it with the arc length *s* such that that *z*(*s*) = *x*(*s*) + *iy*(*s*) describes *∂*Ω in the anticlockwise direction with 0 ≤ *s* ≤ *L*.

The unit normal and tangent vectors on *∂*Ω are then given by:

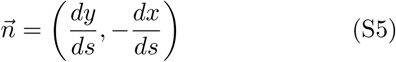

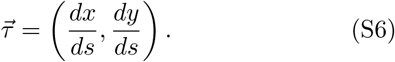

To reformulate Eqn. S4 in terms of a complex contour integral, we define the streamline function 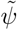 such that 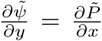 and 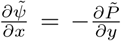 (and hence 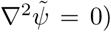). The complex potential 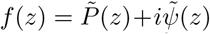 is then an analytic function, and so is its derivative 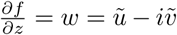, where *ũ* and 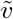 are respectively the *x* and *y* components of **ũ**.

We then define the function 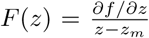, such that *F* (*z*) is a analytic function inside the tissue except at the simple pole *z* = *z*_*m*_. If the pole *z*_*m*_ lies on the boundary, based on Cauchy’s integral formula,

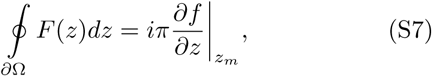

which can be rewritten as:

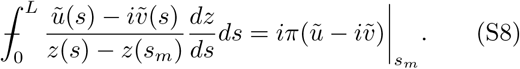

Since 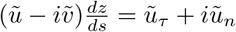, this can be expressed as:

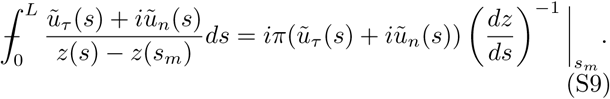

since *P* = 0 on *∂*Ω, *ũ*_*τ*_ = −*û*_*τ*_ on the tissue boundary. We therefore have

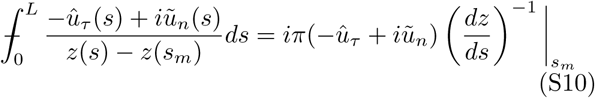

for any 0 ≤ *s*_*m*_ ≤ *L*, and *û*_*τ*_ (*s*) known from Eqn. S3.

By discretizing the boundary and setting *s*_*m*_ to be the mid-points of the mesh, these set of equations can be solved numerically to obtain *ũ*_*n*_ at each of the discrete mesh points. The total velocity is then found by adding this to *û*_*n*_ (which is also known from Eqn. S3).

## Appendix B: Initialization with an ellipse

Initialization with an elliptic (instead of parabolic) cluster of the same dimensions can give qualitatively similar structural features.

**FIG. S1.**
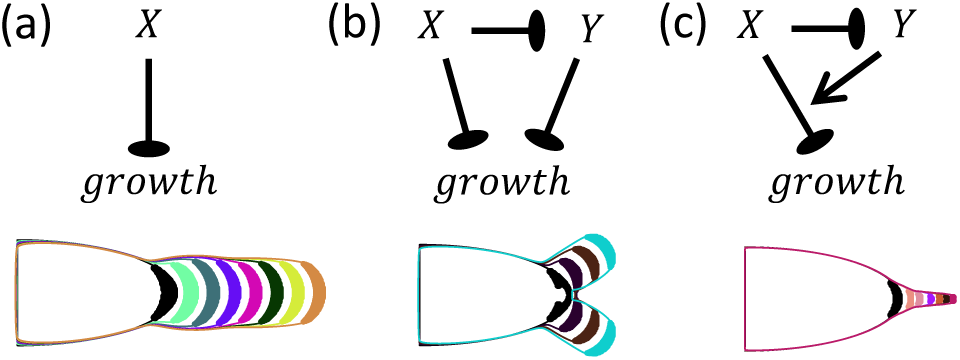
Growth dynamics with an initial elliptic cluster for the following regulatory schemes: (a) Single growth inhibitor [parameters: 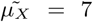], (b) 2 growth inhibitors [parameters: 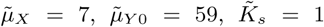, *γ*_*r*_ *=* 1], and (c) 1 growth inhibitor and 1 growth-threshold regulator [parameters: 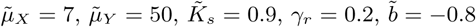]. [Other parameters: initial tissue length 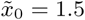, initial tissue width 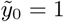]

## Notes

### Competing Interest Statement

The authors have declared no competing interest.

